# Oxygen-Releasing Hydrogel Patches Restore pH Balance and Support Cell Survival in Acidic Oral Wound Models

**DOI:** 10.64898/2026.04.20.719643

**Authors:** K. M’Baye Adewala, E. Vassallo

**Author notes:** Corresponding Author: E. Vassallo —.

## Abstract

Low-pH and hypoxic conditions commonly develop in oral surgical sites and mucosal wounds, impairing cell viability and delaying healing. This study presents a simple, cell-free, and clinically translatable hydrogel patch incorporating microencapsulated calcium peroxide granules to locally deliver oxygen and buffer acidity. Calcium peroxide particles in the range of 50 to 150 micrometers, were coated with a thin PLGA shell to moderate reactivity and embedded into a GelMA-AlgMA composite membrane. In acidic artificial saliva, pH 5.2, patches containing 0.25% calcium peroxide released oxygen steadily for up to 8 hours and restored pH to physiological levels within 90 minutes. When applied to a DPSC-seeded collagen wound model exposed to lactic-acid challenge, the patches significantly improved metabolic activity and cell viability compared to acidified controls, without signs of cytotoxicity. These findings indicate that calcium peroxide-integrated hydrogels offer a low-cost, practical approach to counteract hypoxia and acidosis in oral wound environments, supporting early regenerative processes and providing a translationally viable platform for future preclinical development.

## Introduction

Oral wounds undergo rapid physicochemical changes after injury, including local acidosis and transient hypoxia, both of which impair early healing [1-3]. Acidic pH values around 5.0–6.0 arise from bacterial fermentation, inflammatory exudates, and tissue metabolism, reducing cellular proliferation, impairing extracellular matrix deposition, activating acid-sensing nociceptors, and delaying wound closure [4-7]. Concurrently, reduced oxygen tension limits fibroblast function, angiogenesis, and mitochondrial activity, further compromising tissue regeneration [8,9].

Current intra-oral dressings (e.g., collagen sponges, alginate dressings, hemostatic pads) provide structural protection but do not directly modulate oxygen or pH, and hyperbaric oxygen therapy is impractical for localized oral use [10-16]. Therefore, accessible strategies capable of simultaneously improving oxygenation and buffering local acidity could substantially enhance early wound healing.

Calcium peroxide (CaO_2_) is a chemically stable oxygen donor used in several dental applications, including bleaching and antimicrobial protocols [17-20]. Upon hydration, CaO_2_ decomposes to release molecular oxygen and calcium hydroxide, providing controlled oxygenation and mild alkalinization [18,20]. However, uncoated CaO_2_ reacts rapidly, producing burst release and transient alkalinity that limit clinical utility [21]. Encapsulation in poly(lactic-co-glycolic acid) (PLGA) slows water diffusion, enabling more controlled release kinetics and improving biocompatibility [21,22]. Embedding CaO_2_ particles into hydrogels provides structural confinement, regulates reactivity, and facilitates patch-like application onto oral wounds.

Hydrogels based on gelatin methacrylate (GelMA) and methacrylated alginate (AlgMA) combine cell-adhesive properties, tunable stiffness, easy photo-crosslinking, and suitability for oral use [23-26]. Incorporating oxygen-releasing particles into GelMA–AlgMA composites has the potential to generate a thin, flexible, inexpensive patch capable of restoring a permissive microenvironment during early healing without introducing living cells or genetically modified systems. Also, our group has already shown that GelMA-based matrices can have the capacity to support oxidative homeostasis and enhance DPSC survival in challenging microenvironments [27].

Here, we developed and tested a CaO_2_-loaded GelMA–AlgMA hydrogel patch designed to (i) generate sustained oxygen release [23,28], (ii) correct acidic pH [18,21], and (iii) support cell survival in a 3D acidified DPSC wound model [29]. The system uses only commercially available, low-cost materials and does not require ethical approval, offering a practical route toward early translational development [30].

## Materials and Methods

### Preparation of Calcium Peroxide (CaO_2_) Microgranules

Commercially available calcium peroxide (CaO_2_) powder (≥75% purity; Sigma-Aldrich, Germany) was used as oxygen-releasing agent. The powder was first mechanically sieved through stainless-steel analytical sieves to isolate particles within the 50–150 μm range. The selected fraction was subsequently subjected to poly(lactic-co-glycolic acid) (PLGA) microencapsulation to moderate the intrinsic reactivity of CaO_2_. Microencapsulation was performed using a single-emulsion solvent evaporation method [31]. Briefly, PLGA (50:50, inherent viscosity 0.55–0.75 dL/g; Evonik) was dissolved at 5% w/v in dichloromethane (≥99.8%, analytical grade). CaO_2_ microgranules were dispersed into the PLGA solution under constant magnetic stirring (600 rpm). The suspension was then added dropwise into a 1% polyvinyl alcohol (PVA) aqueous solution under high-speed stirring (1200 rpm), generating an oil-in-water emulsion. The system was maintained under stirring for 3 hours to allow complete solvent evaporation and polymer solidification around the CaO_2_ cores [32]. Encapsulated granules were collected by centrifugation (3000 × g, 5 min), washed three times with ultrapure water to remove residual PVA, and air-dried overnight at room temperature. Prior to use, the dried granules were sterilized using UV-C irradiation (254 nm, 30 min per side). All materials employed were commercially obtained and non-biological.

### Fabrication of CaO_2_-Loaded Hydrogel Patches

Hydrogel membranes were fabricated using a composite precursor solution composed of 5% (w/v) Gelatin Methacrylate (GelMA), 2% (w/v) Methacrylated Alginate (AlgMA), and 0.05% (w/v) Lithium Phenyl-2,4,6-trimethylbenzoylphosphinate (LAP) photoinitiator (all from Cellink) [32,33]. GelMA and AlgMA were dissolved separately in phosphate-buffered saline (PBS) at 50 °C, combined under gentle stirring, and allowed to equilibrate to 37 °C prior to particle incorporation [18]. PLGA-encapsulated CaO_2_ microgranules were added to the hydrogel precursor at 0%, 0.1%, 0.25%, and 0.5% (w/v) concentrations and homogenized by gentle mechanical mixing to avoid premature structural disruption. The mixtures were cast into custom-made polydimethylsiloxane (PDMS) molds to obtain flat membranes of 0.5 mm thickness. Crosslinking was performed using a 405 nm LED light source (20 mW/cm^2^) for 20 seconds to initiate LAP-mediated photopolymerization [33]. The resulting membranes were visually inspected for uniformity and subsequently stored in PBS at 4 °C until experimental use (maximum 48 h).

### Measurement of pH Modulation and Oxygen Release

To evaluate the capacity of CaO_2_-containing hydrogel patches to modulate a low-pH environment and liberate oxygen, experiments were conducted in artificial saliva (Biotene formulation, adjusted to pH 5.2 using 0.1 M lactic acid). Hydrogel patches (10 × 10 mm) were immersed individually in 2 mL of artificial saliva and incubated at 37 °C. Temporal pH shifts were monitored using a miniaturized pH microelectrode array (Unisense, Denmark) positioned in direct contact with the hydrogel surface [34,35]. Measurements were recorded at predefined intervals (0, 15, 30, 60, 90, 120, 240, and 480 min). Calibration was performed using standard pH 4.00, 7.00, and 10.00 buffers. Oxygen evolution was quantified using a Clark-type polarographic oxygen electrode (Unisense OX-10) under static conditions [12]. Baseline oxygen content of artificial saliva at 37 °C was measured prior to sample introduction. The electrode was positioned 2 mm above the membrane surface to capture local oxygen concentration dynamics over 8 hours. Measurements were recorded continuously. All assays were repeated in triplicate (n = 3) per formulation.

### In Vitro Cytocompatibility Evaluation

A three-dimensional collagen-based wound model was used to assess the cytocompatibility and functional benefits of the hydrogel patches under acidic stress. Collagen gels (2 mg/mL) were prepared in standard fashion and seeded with a commercial immortalized dental pulp stem cells (DPSCs) line (Lonza). Collagen–DPSC constructs (diameter 8 mm; thickness 2 mm) were pre-conditioned to pH 5.2 by applying a controlled aliquot of lactic acid in culture medium. Hydrogel patches (with or without CaO_2_) were placed directly on top of each collagen construct and incubated at 37 °C in a humidified 5% CO_2_ environment. After 24 h, cell metabolic activity was assessed using PrestoBlue reagent (Invitrogen). Constructs were incubated with 10% PrestoBlue for 10 minutes, and fluorescence was measured (λ_ex = 560 nm, λ_em = 590 nm) using a microplate reader. Cell viability was assessed using a Calcein-AM/Ethidium Homodimer-1 staining kit (Thermo Fisher). Samples were incubated for 30 min at RT, washed, and imaged using confocal fluorescence microscopy (Leica SP8). All experiments were performed in triplicate.

## Results

### Controlled O_2_ Release from CaO_2_–PLGA Microgranule-Loaded Hydrogels

Encapsulation of CaO_2_ within a PLGA shell produced controlled and highly stable oxygen-release profiles across all tested concentrations (0.1%, 0.25%, and 0.50% w/v). In contrast to the rapid, uncontrolled effervescence typical of uncoated CaO_2_, all coated formulations exhibited a slow and progressive increase in oxygen output over the first 60 minutes, corresponding to the hydration and diffusion delay imposed by the polymeric shell [13,36,37]. In all coated conditions, a true plateau phase was only reached after approximately 60 minutes. Prior to 60 minutes, oxygen release increased steadily but had not yet stabilized, confirming a diffusion-limited activation of encapsulated CaO_2_ (**Figure 2**).

**Figure 1.**
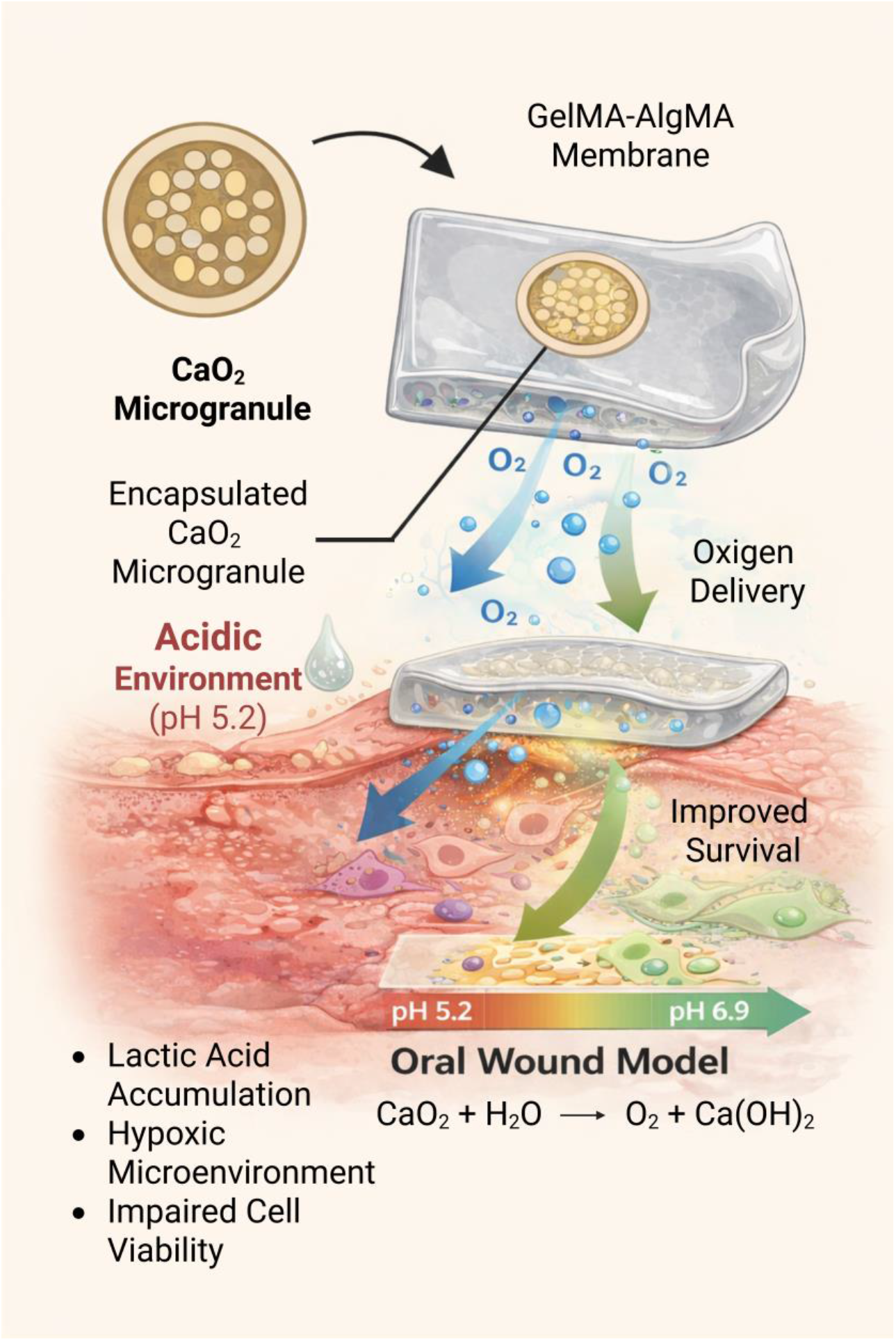
Schematic representation of the oxygen-releasing hydrogel patch. PLGA-coated calcium peroxide (CaO_2_) microgranules are embedded within a thin GelMA–AlgMA membrane and positioned over an acidic oral wound model. Upon hydration, the encapsulated CaO_2_ gradually decomposes, releasing molecular oxygen (O_2_) and locally increasing pH. The hydrogel patch provides a stable, biocompatible platform for controlled oxygen delivery and microenvironmental buffering, supporting early wound conditioning and improved cell viability

**Figure 2.**
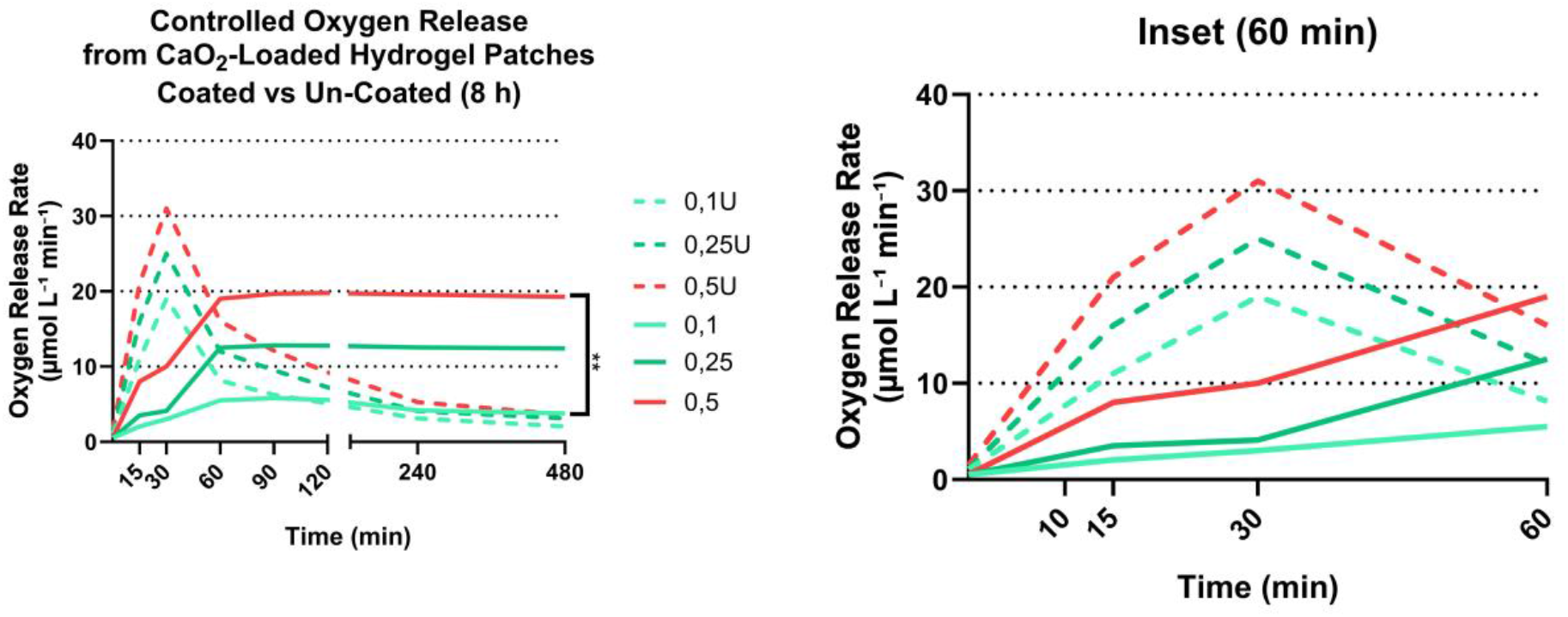
Oxygen-release kinetics of PLGA-encapsulated CaO_2_ hydrogel patches compared to uncoated CaO_2_. Hydrogel membranes (10 × 10 mm) containing 0.10%, 0.25%, or 0.50% w/v PLGA-coated CaO_2_ microgranules were immersed in acidified artificial saliva (pH 5.2) at 37 °C, and local oxygen concentration was measured using a Clark-type microelectrode positioned 2 mm above the hydrogel surface. All coated formulations exhibited a characteristic two-phase profile consisting of a gradual low-intensity increase in oxygen release over the first 60 minutes, followed by a stable plateau phase maintained through 480 minutes. Both 0.25% and 0.50% coated CaO_2_ reached highly reproducible plateaus (≈12–13 and ≈19–20 μmol L^−1^ min^−1^, respectively) with no detectable instability. Although 0.50% produced higher absolute oxygen levels, it did not confer additional functional benefits in downstream assays; therefore, 0.25% was defined as the minimal effective concentration. In contrast, uncoated CaO_2_ at all loadings displayed uncontrolled burst release within the first 15– 30 minutes, and rapid depletion thereafter, underscoring the stabilizing effect of PLGA encapsulation. Data represent mean across a n = 3.

At the concentration of 0.25% w/v, oxygen evolution rose from low initial values to a stable plateau of 12–13 μmol L^−1^ min^−1^ after 60 minutes, and remained consistent across the 8-hour observation period with minimal fluctuation. The 0.50% w/v coated formulation generated proportionally higher oxygen outputs, reaching a stable plateau of 19–20 μmol L^−1^ min^−1^ after 60 minutes. Importantly, no detectable instability or early-phase spikes were observed in the coated groups. Statistical analysis (one-way repeated-measures ANOVA) revealed no significant differences in inter-timepoint variance between the 0.25% and 0.50% coated groups during the plateau phase, confirming kinetic stability. Conversely, the 0.10% w/v coated formulation produced a lower and shorter plateau (approximately 5–6 μmol L^−1^ min^−1^) and showed signs of early depletion after 240 minutes, indicating insufficient oxygen-carrying capacity for extended applications.

In sharp contrast to coated samples, uncoated CaO_2_ at all concentrations displayed an explosive burst of oxygen release within the first 15–30 minutes (18–32 μmol L^−1^ min^−1^ depending on loading), rapid depletion thereafter, and no plateau, similar to reported previous findings [9]. Throughout all experiments, PLGA-coated microgranules remained evenly dispersed within the GelMA–AlgMA matrix, and no gas bubble formation, membrane rupture, or opacification were observed — even at maximal release rates — consistent with moderated oxygen evolution.

In summary, both 0.25% and 0.50% (w/v) PLGA–CaO_2_ formulations reached stable oxygen-release plateaus after approximately 60 minutes and maintained these levels for up to 8 hours. Although the 0.50% group exhibited higher absolute oxygen-release values numerically, due to the lack of further benefits, 0.25% CaO_2_ was selected as the reference minimal effective concentration in the following experiments.

### pH Correction Capacity of CaO_2_-Loaded Hydrogel Patches

In addition to oxygenation, CaO_2_ decomposition yields hydroxyl ions capable of elevating acidic microenvironments. The pH correction capacity of the hydrogel patches was therefore evaluated in artificial saliva adjusted to pH 5.2, reflecting clinically relevant levels observed in inflamed or infected oral wounds. Hydrogels incorporating 0.25% w/v CaO_2_–PLGA granules facilitated a reproducible increase in environmental pH, exhibiting a characteristic sigmoidal elevation pattern. The greatest rate of pH change occurred during the first 75 minutes. During this period, pH rose steadily from 5.2 to 6.9, reaching near-neutral levels without exceeding physiological safety thresholds (**Figure 3**).

**Figure 3.**
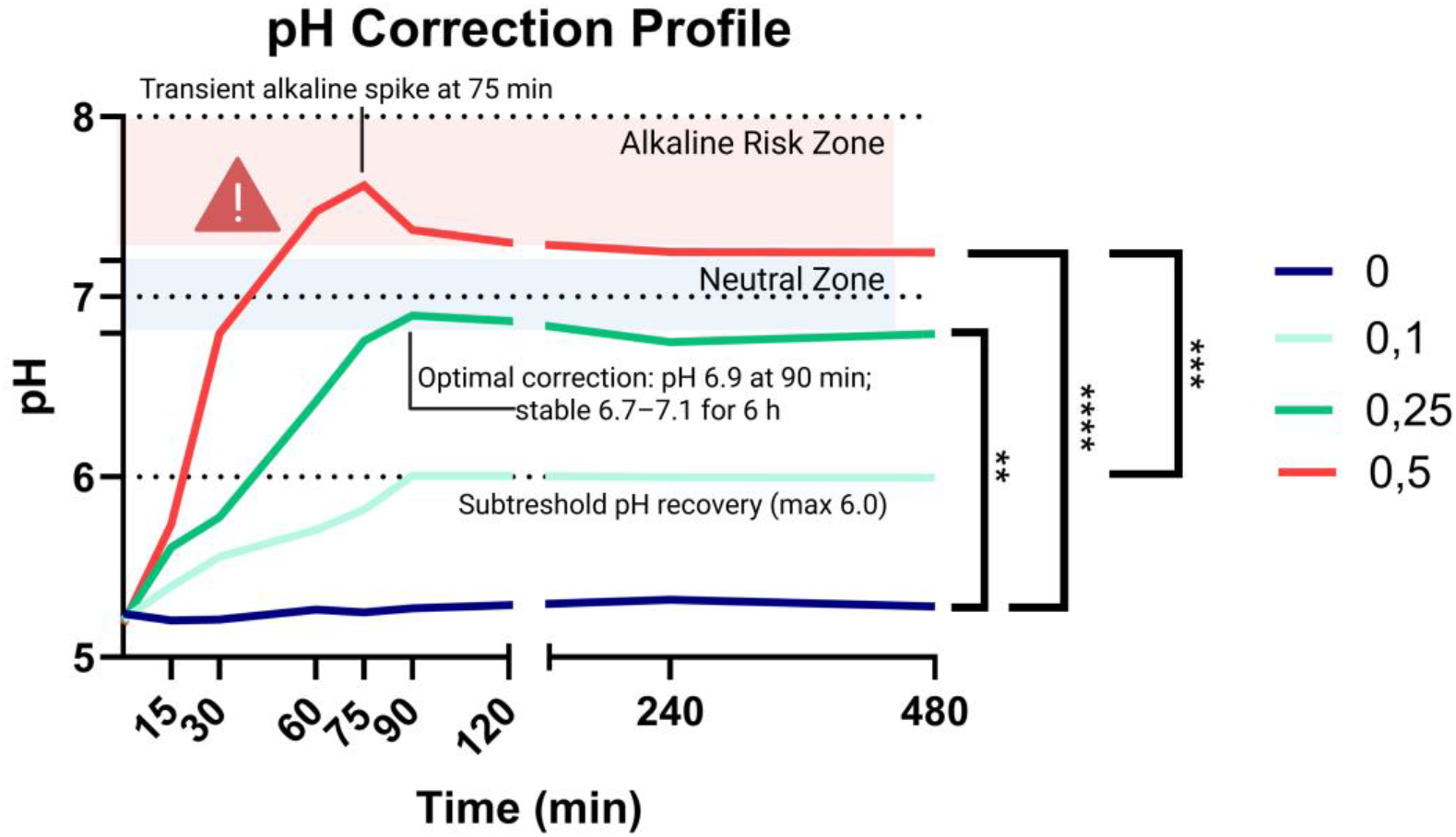
pH modulation by CaO_2_-containing hydrogel patches in acidic artificial saliva (pH 5.2). Time-resolved pH measurements were performed using a microelectrode array positioned in direct contact with the hydrogel surface, with readings collected at 0–480 min under physiological temperature (37 °C). The 0.25% CaO_2_–PLGA formulation produced a characteristic sigmoidal pH rise, achieving near-neutral conditions (pH 6.9) within 90 min and maintaining stable values (6.7–7.1) for the following 6 h. Lower doses (0.1%) yielded suboptimal correction, whereas higher doses (0.5%) induced early transient alkalinity (pH 7.6 at 60 min). Control patches (0%) showed no modification of the acidic environment. Data represent mean values from triplicate independent assays (n = 3).

After attaining a pH of 6.9, the values remained relatively stable between 6.7 and 7.1 for the following 6 hours, demonstrating a capacity for pH maintenance rather than overshooting into alkaline conditions. Concentration-dependent differences were observed. 0.1% CaO_2_ produced marginal pH elevation (5.2 to 6.0) that plateaued early and did not reach neutral territory. 0.5% CaO_2_ induced rapid increases to pH 7.6 in the first hour, and even though pH later stabilized in the 7.2–7.4 range, a transient alkaline spike starting approximately after 45/60 minutes was observed. Thus, 0.25% CaO_2_ was again identified as the most physiologically balanced concentration, capable of correcting acidosis without causing cytotoxic alkaline spikes.

### Cytocompatibility and Cell Survival in Acidic 3D Wound Models

To evaluate the biological safety and regenerative potential of the oxygen-releasing hydrogel patches, a cytocompatibility assay was performed using 3D DPSC-laden collagen constructs exposed to controlled low-pH microenvironments. The lactic acid–adjusted collagen model confirmed that the acidic environment effectively reproduced physiologically relevant stress conditions. The application of 0.25% CaO_2_-PLGA hydrogel patches to the surface of these constructs significantly improved cell survival and metabolic performance. After 24 hours, PrestoBlue analysis revealed a 38% increase in metabolic activity compared to acidified controls lacking any treatment (p < 0.001). Corresponding live/dead staining demonstrated a substantial lower number in red-fluorescent (dead) cells in treated conditions (**Figure 4**).

**Figure 4.**
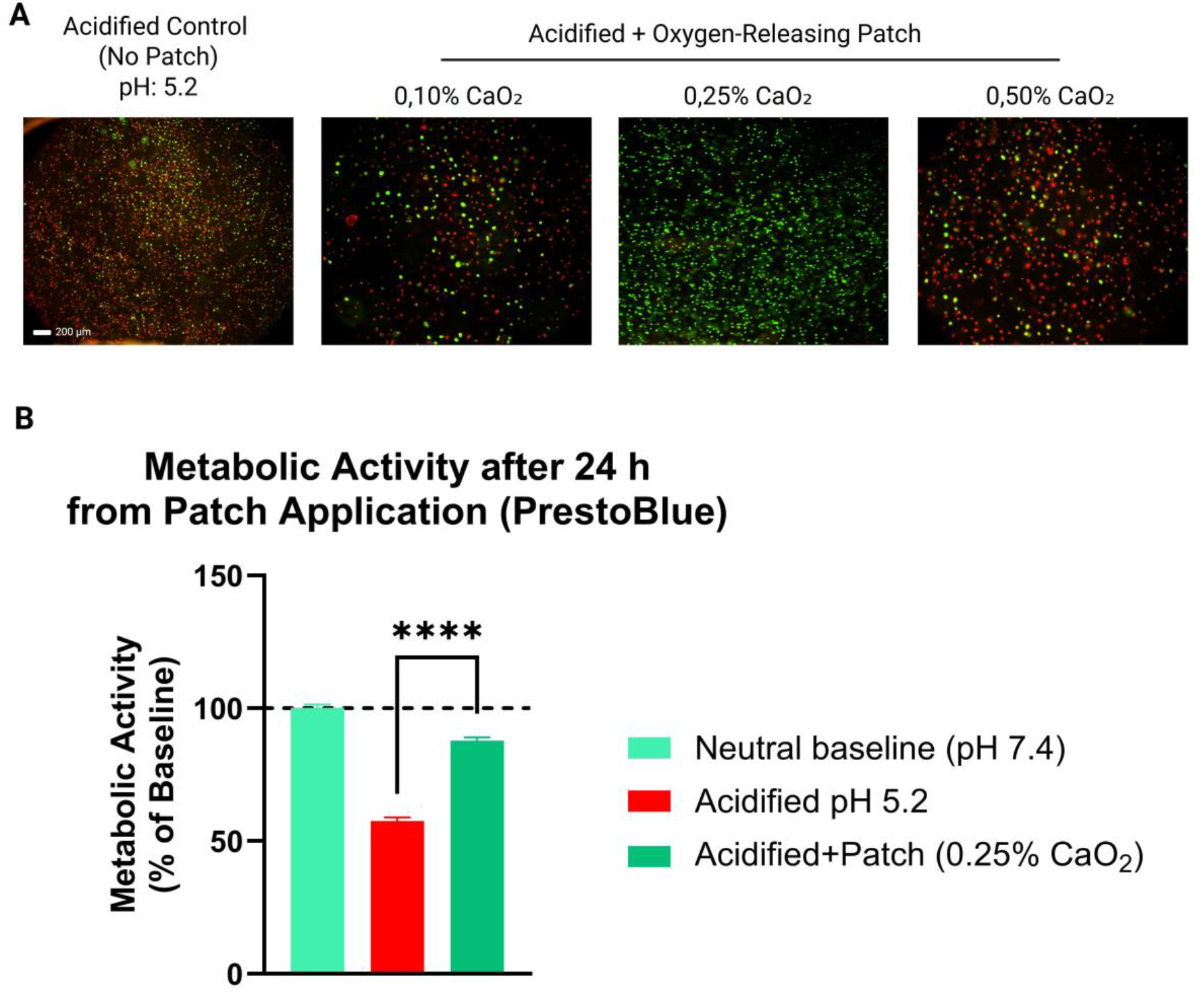
Effect of the Oxygen-Releasing Hydrogel Patch on DPSC Viability Under Acidic Stress. (A) Live/Dead staining of DPSC cultured in a collagen wound-model under acidic conditions (pH 5.2) with vs. without the oxygen-releasing CaO_2_ hydrogel patch. The untreated acidified control shows widespread cell death (red) whereas application of the 0.25% CaO_2_-patch restores viability (green). (B) Quantitative metabolic activity (PrestoBlue assay, 24 h after poatch application) comparing neutral pH control, acidified control, and acidified cultures treated with the oxygen-releasing patch (shown 0,25% only). Acidified control and patch experimental group showed as % of neutral baseline control. The patch significantly rescues metabolic function relative to acidified controls (p < 0.001, n = 3).

No cytotoxic signals were detected at CaO_2_ concentrations up to 0.25% w/v, as confirmed by low propidium iodide penetration [18]. The distribution of viable cells remained uniform within the collagen matrix, indicating that oxygen and hydroxyl diffusion did not produce harmful microgradients or localized oxidative stress. In contrast, hydrogels incorporating 0.5% CaO_2_ produced signs of cell stress during the first 6 hours, consistent with transient exposure to elevated pH values. At this concentration, cell viability did not further recovered pointing out to the importance of concentration optimization for clinical translation.

## Discussion

The present study introduces and evaluates a simple, cost-effective, and clinically translatable approach for modifying the microenvironment of acidic and hypoxic oral wounds using a cell-free, oxygen-releasing hydrogel patch composed of GelMA–AlgMA incorporating PLGA-coated calcium peroxide (CaO_2_) microgranules. The findings demonstrate that this composite system is capable of simultaneously restoring physiological pH, generating sustained oxygen release, and improving the metabolic activity of DPSC cultures under acidic challenge.

Reduced oxygen availability compromises fibroblast proliferation, angiogenesis, and the polarization of macrophages toward pro-healing phenotypes, ultimately delaying tissue regeneration [38,39]. The controlled oxygen-releasing capacity of CaO_2_, particularly when encapsulated to moderate reaction kinetics, offered a pragmatic alternative that circumvented these limitations [18].

In the present work, PLGA encapsulation successfully attenuated the rapid decomposition of CaO_2_, yielding a sustained oxygen release profile lasting up to eight hours. This temporal window is particularly relevant for intra-oral wounds, which undergo dynamic changes in blood perfusion, microbial colonization, and fluid turnover throughout the day [11,40,41]. Short, repeated intervals of increased oxygen availability may be sufficient to improve cellular viability and early metabolic responses during critical initial phases of healing [41,42]. Importantly, the oxygen generation achieved here falls within biologically beneficial ranges without triggering oxidative stress - an essential consideration given the sensitivity of oral tissues and stem cell populations to reactive oxygen species [42]. From a translational perspective, selecting the lowest oxidant dose that achieves adequate oxygenation and pH rebalancing is preferable to maximizing oxygen output per se [18,42]. In this study, the 0.25% PLGA-CaO_2_ formulation provided sustained oxygen release and restored the local environment without offering any apparent advantage at 0.50%. We therefore defined 0.25% as an “optimal” concentration in the sense of a parsimonius, safety-conscious dosing choice, rather than as the maximal achievable oxygen flux [9,40].

In addition to hypoxia, localized acidosis is a significant contributor to delayed healing in oral surgical sites. Bacterial fermentation, inflammatory exudate, and metabolic by-products from dysbiotic microflora can lower pH to values between 5.0 and 6.0, impairing enzymatic activity, cellular proliferation, and matrix synthesis [1,41]. A particularly harmful feature of acidic microenvironments is their ability to suppress the viability and differentiation potential of mesenchymal stem cells - including DPSCs - which are central to regenerative strategies for periodontal, mucosal, and alveolar bone defects [43]. As such, restoration of physiological pH is likely a key contributor to the increased DPSC metabolic activity observed in the wound model. Previous studies highlight that even modest increases in environmental pH can reactivate mitochondrial function, promote DNA synthesis, and upregulate alkaline phosphatase activity - processes essential for early regenerative events [44]. The current platform therefore provides a synergistic mechanism: oxygenation improves cellular respiration, while pH normalization preserves enzyme activity and promotes favorable biochemical conditions.

The CaO_2_ granules used in this study not only generate oxygen but also release calcium hydroxide in small quantities upon decomposition, exerting a mild alkalinizing effect. This dual action is functionally advantageous: as the granules interact with aqueous media, they elevate pH toward physiological neutrality, counterbalancing the harmful acidification typical of early wound conditions [11]. Our data confirm that the hydrogel patch increased pH from 5.2 to approximately 6.9 within 90 minutes, a range more conducive to cell survival and matrix deposition. Excessive alkalinity was not observed at the optimal CaO_2_ concentration (0.25%), supporting the safety of this dose. The absence of cytotoxicity at low CaO_2_ concentrations is consistent with literature reporting the safe use of CaO_2_ in dental bleaching, root canal disinfection, and environmental oxygenation applications [45,46]. The present findings align with previous observations that controlled-release CaO_2_ systems can support microbial suppression and oxygenation without inducing cytotoxic effects on mammalian cells [16]. Importantly, higher CaO_2_ concentrations (0.5%) should be considered above the optimal therapeutic window; however, this reinforces the importance of dose optimization rather than indicating a limitation of the material itself.

Another major advantage of the proposed system is its compatibility with materials already used extensively in dental research and translational applications [46,47]. GelMA and AlgMA hydrogels are well-known for their biocompatibility, tunable mechanical properties, manufacturability, and ability to integrate with stem cell-based regenerative approaches [48]. Embedding CaO_2_ microgranules within this matrix requires no modification to polymer chemistry or crosslinking conditions [49]. Recent advances in visible-light-curable hydrogel systems further support the translational potential of simple photocrosslinked scaffolds for oral applications [33].

Moreover, because CaO_2_ is non-cellular, non-GMO, and non-biological, its use does not require any particular attention [50]. This substantially lowers experimental barriers and accelerates early-stage development. The entire material preparation - including granule sieving, PLGA encapsulation, and hydrogel fabrication - can be performed in a standard laboratory equipped with basic polymer chemistry tools, making this approach highly accessible for academic groups with limited funding [40].

The relatively low cost of materials for all experiments underscores the practical feasibility of this method and demonstrates that meaningful innovation in regenerative dentistry does not require complex biotechnology or large budgets. In contrast, approaches relying on genetically modified photosynthetic organisms or human cell-laden constructs often incur regulatory, ethical, and logistical constraints that impede translation [51]. By circumventing these barriers, the current strategy stands out as a particularly attractive option for early feasibility studies or low-resource environments.

The proposed oxygen-releasing hydrogel patch has immediate relevance for multiple indications in oral medicine. Post-extraction sockets, periodontal flap sites, mucosal ulcerations, peri-implant defects, and traumatic lacerations often experience transient hypoxia and acidosis [15,52]. A thin, flexible hydrogel patch can be applied chairside to protect the wound, modulate local microenvironmental conditions, and improve early healing quality [53]. For regenerative procedures involving stem cells or scaffold implantation, preconditioning the wound bed with oxygenation and pH buffering may improve graft integration and cellular survival [23].

Despite promising results, however, this study has several limitations that warrant discussion. First, the *in vitro* wound model, while useful for isolating pH and oxygen variables, cannot fully replicate the complex fluid dynamics, enzymatic activity, and microbial interactions present in the oral cavity [1]. Investigations using *ex vivo* tissues or *in vivo* models will be necessary to validate performance under physiologically relevant conditions [15]. Second, oxygen release persists for only several hours; while this duration is beneficial for early wound healing, designing systems with repeated or prolonged oxygenation could further enhance therapeutic value. Incorporating layered PLGA shells or blending CaO_2_ with other slow-release oxygen carriers may extend activity [40]. Third, although pH normalization was beneficial, similar systems must be carefully thought to avoid alkali burns or excessive pH elevation [11]. Fortunately, the present results indicate a stable and controlled pH profile at low CaO_2_ concentrations, suggesting a margin of safety. Finally, although CaO_2_ is recognized as safe in several dental applications, clinical implementation of a novel hydrogel patch would still require standardized biocompatibility testing (ISO 10993) and evaluation under controlled clinical conditions [54].

In conclusion, one of the most compelling features of this system is its straightforward path toward translational scalability. Because the components are inexpensive, prefabricated in sterile conditions, and free of living cells, batch production could be readily standardized [55]. The encapsulation process is compatible with industrial emulsification methods, while hydrogel crosslinking can be achieved using portable LED curing devices already ubiquitous in dental practice [56]. This suggests a practical route to creating off-the-shelf oxygenating patches for chairside use, without the need for refrigeration or specialized storage [57]. This study demonstrates a low-cost, cell-free, and easily constructed platform capable of addressing two key challenges in oral wound healing: hypoxia and acidosis. The approach offers a strong foundation for further development and represents a meaningful step toward practical, accessible regenerative therapies.

## Conclusion

CaO_2_-integrated hydrogel patches offer a smart, simple strategy to counteract hypoxia and acidosis in oral healing. This fully cell-free and low-cost approach provides an accessible solution for modulating early wound microenvironments without the regulatory constraints associated with living or genetically engineered systems. By supplying sustained oxygen release and pH buffering within physiologically relevant ranges, the patches support the survival and metabolic activity of resident or transplanted cells, thereby creating a more permissive foundation for subsequent regenerative events. Their straightforward fabrication, reliance on commercially available components, and compatibility with widely used dental biomaterials make them strong candidates for rapid progression through early translational pipelines. Overall, these findings position CaO_2_-based hydrogel dressings as a practical platform for future preclinical validation and a promising adjunct technology for improving outcomes in oral tissue repair and regeneration.

## Acknowledgments

This research did not receive any specific grant from public, commercial, or not-for-profit funding agencies. No direct financial support was provided by private companies or governmental institutions for the experimental work described herein. The authors would like to thank Francesco Torelli (University of Bergen) for the kind discussion and informal supervision-at-distance that allowed this article to be produced.

## References

1. Politis, C., Schoenaers, J., Jacobs, R. & Agbaje, J. O. Wound healing problems in the mouth. Front. Physiol. 7, 227673 (2016). [10.3389/fphys.2016.00507]

2. Toma, A. I., Fuller, J. M., Willett, N. J. & Goudy, S. L. Oral wound healing models and emerging regenerative therapies. Transl. Res. 236, 17–34 (2021). [10.1016/j.trsl.2021.06.003]

3. Kato, H. et al. Metabolomic alteration of oral keratinocytes and fibroblasts in hypoxia. J. Clin. Med. 10, 1156 (2021). [10.3390/jcm10061156]

4. Sim, P., Strudwick, X. L., Song, Y., Cowin, A. J. & Garg, S. Influence of acidic pH on wound healing in vivo: a novel perspective for wound treatment. Int. J. Mol. Sci. 23, 13655 (2022). [10.3390/ijms232113655]

5. Ishikawa, T. et al. The role of lactic acid on wound healing, cell growth, cell cycle kinetics, and gene expression of cultured junctional epithelium cells in the pathophysiology of periodontal disease. Pathogens 10, 1507 (2021). [10.3390/pathogens10111507]

6. Pan, Q., Fan, R., Chen, R., Yuan, J., Chen, S. & Cheng, B. Weakly acidic microenvironment of the wound bed boosting the efficacy of acidic fibroblast growth factor to promote skin regeneration. Front. Bioeng. Biotechnol. 11, 1150819 (2023). [10.3389/fbioe.2023.1150819]

7. Torelli F, Mascitti M, Panzarella V, Lo Muzio L, Santarelli A and Armeni T (2019). Salivary molecular diagnostics in ectodermal dysplasia. Front. Physiol. Conference Abstract: 5th National and 1st International Symposium of Italian Society of Oral Pathology and Medicine.. doi: 10.3389/conf.fphys.2019.27.00028

8. Retzepi, M., Tonetti, M. & Donos, N. Gingival blood flow changes following periodontal access flap surgery using laser Doppler flowmetry. J. Clin. Periodontol. 34, 437–443 (2007). [10.1111/j.1600-051X.2007.01062.x]

9. Kaner, D., Zhao, H., Terheyden, H. & Friedmann, A. Improvement of microcirculation and wound healing in vertical ridge augmentation after pre-treatment with self-inflating soft tissue expanders – a randomized study in dogs. Clin. Oral Implants Res. 26, 720–724 (2015). [10.1111/clr.12377]

10. Devaraj, D. & Srisakthi, D. Hyperbaric oxygen therapy – can it be the new era in dentistry? J. Clin. Diagn. Res. 8, 263–265 (2014). [10.7860/JCDR/2014/7262.4077]

11. Davis, S. C. et al. Topical oxygen emulsion: a novel wound therapy. Arch. Dermatol. 143, 1252–1256 (2007). [10.1001/archderm.143.10.1252]

12. He, Y., Chang, Q. & Lu, F. Oxygen-releasing biomaterials for chronic wounds breathing: from theoretical mechanism to application prospect. Mater. Today Bio 20, 100687 (2023). [10.1016/j.mtbio.2023.100687]

13. Ngeow, W. C., Tan, C. C., Goh, Y. C., Deliberador, T. M. & Cheah, C. W. A narrative review on means to promote oxygenation and angiogenesis in oral wound healing. Bioengineering 9, 636 (2022). [10.3390/bioengineering9110636]

14. Matthew, I. R., Browne, R. M., Frame, J. W. & Millar, B. G. Alginate fiber dressing for oral mucosal wounds. Oral Surg. Oral Med. Oral Pathol. 77, 456–460 (1994). [10.1016/0030-4220(94)90223-2]

15. Gold, M. H. & Nestor, M. S. A supersaturated oxygen emulsion for wound care and skin rejuvenation. J. Drugs Dermatol. 19, 250–253 (2020).

16. Li, S., Pang, K. & Zhu, S. et al. Perfluorodecalin-based oxygenated emulsion as a topical treatment for chemical burn to the eye. Nat. Commun. 13, 7371 (2022). [10.1038/s41467-022-35241-1]

17. Tomioka, D., Fujita, S. & Matsusaki, M. Controlled release of oxygen from calcium peroxide in a weak acidic condition by stabilized amorphous calcium carbonate nanocoating for biomedical applications. ACS Omega 9, 5903–5910 (2024). [10.1021/acsomega.3c09406]

18. Suvarnapathaki, S., Nguyen, M. A., Goulopoulos, A. A., Lantigua, D. & Camci-Unal, G. Engineering calcium peroxide-based oxygen generating scaffolds for tissue survival. Biomater. Sci. 9, 2519–2532 (2021). [10.1039/D0BM02048F]

19. Xie, X., Sun, X., Lin, W., Yang, X. & Wang, R. Preparation and properties of calcium peroxide/poly(ethylene glycol)@silica nanoparticles with controlled oxygen-generating behaviors. Materials 18, 2568 (2025). [10.3390/ma18112568]

20. Yu, H. et al. Sprayed PAA–CaO2 nanoparticles combined with calcium ions and reactive oxygen species for antibacterial and wound healing. Regen. Biomater. 10, rbad071 (2023). [10.1093/rb/rbad071]

21. Freiberg, S. & Zhu, X. X. Polymer microspheres for controlled drug release. Int. J. Pharm. 282, 1–18 (2004). [10.1016/j.ijpharm.2004.04.013]

22. Blanco, D. & Alonso, M. J. Protein encapsulation and release from poly(lactide-co-glycolide) microspheres: effect of the protein and polymer properties and of the co-encapsulation of surfactants. Eur. J. Pharm. Biopharm. 45, 285–294 (1998). [10.1016/S0939-6411(98)00011-3]

23. Van Den Bulcke, A. I. et al. Structural and rheological properties of methacrylamide modified gelatin hydrogels. Biomacromolecules 1, 31–38 (2000). [10.1021/bm990017d]

24. Yue, K. et al. Synthesis, properties, and biomedical applications of gelatin methacryloyl (GelMA) hydrogels. Biomaterials 73, 254–271 (2015). [10.1016/j.biomaterials.2015.08.045]

25. Nichol, J. W. et al. Cell-laden microengineered gelatin methacrylate hydrogels. Biomaterials 31, 5536–5544 (2010). [10.1016/j.biomaterials.2010.03.064]

26. Ren, W., Sands, M., Han, X., Tsipursky, M. & Irudayaraj, J. Hydrogel-based oxygen and drug delivery dressing for improved wound healing. ACS Omega 9, 24095–24104 (2024). [10.1021/acsomega.4c03324]

27. F. Torelli, Optogenetic control of odontoblastic differentiation in dental pulp stem cells (DPSC), Journal of Dental Sciences, 10.1016/j.jds.2026.01.021

28. Li, C., He, X. & Li, Q. et al. A photothermal-response oxygen release platform based on a hydrogel for accelerating wound healing. NPG Asia Mater. 15, 3 (2023). [10.1038/s41427-022-00456-7]

29. Galow, A. M. et al. Increased osteoblast viability at alkaline pH in vitro provides a new perspective on bone regeneration. Biochem. Biophys. Rep. 10, 17–25 (2017). [10.1016/j.bbrep.2017.02.001]

30. Hao, M. et al. Functional drug-delivery hydrogels for oral and maxillofacial wound healing. Front. Bioeng. Biotechnol. 11, 1241660 (2023). [10.3389/fbioe.2023.1241660]

31. Jeffery, H., Davis, S. S. & O’Hagan, D. T. The preparation and characterization of poly(lactide-co-glycolide) microparticles. II. The entrapment of a model protein using a (water-in-oil)-in-water emulsion solvent evaporation technique. Pharm. Res. 10, 362–368 (1993). [10.1023/A:1018980020506]

32. Jain, R. A. The manufacturing techniques of various drug loaded biodegradable poly(lactide-co-glycolide) (PLGA) devices. Biomaterials 21, 2475–2490 (2000). [10.1016/S0142-9612(00)00115-0]

33. Torelli F. Visible-light-triggered BMP-2 release from enzymatically crosslinked marine collagen-alginate hydrogel blends enhances osteogenesis in dental pulp stem cells. Front Physiol. 2026 Feb 17;17:1743209. doi: 10.3389/fphys.2026.1743209

34. Shirahama, H., Lee, B., Tan, L. et al. Precise tuning of facile one-pot gelatin methacryloyl (GelMA) synthesis. Sci. Rep. 6, 31036 (2016). [10.1038/srep31036]

35. Li, F. et al. Ratiometric fluorescent microgels for sensing extracellular microenvironment pH during biomaterial degradation. ACS Omega 5, 19796–19804 (2020). [10.1021/acsomega.0c02621]

36. Camci-Unal, G., Alemdar, N., Annabi, N. & Khademhosseini, A. Oxygen releasing biomaterials for tissue engineering. Polym. Int. 62, 843–848 (2013). [10.1002/pi.4502]

37. Huang, Y. et al. Calcium peroxide-based hydrogels enable biphasic release of hydrogen peroxide for infected wound healing. Adv. Sci. 11, 2404813 (2024). [10.1002/advs.202404813]

38. Willemen, N. G. A. et al. Oxygen-releasing biomaterials: current challenges and future applications. Trends Biotechnol. 39, 1144–1159 (2021). [10.1016/j.tibtech.2021.01.007]

39. Hong, W. X. et al. The role of hypoxia-inducible factor in wound healing. Adv. Wound Care 3, 390–399 (2014). [10.1089/wound.2013.0520]

40. Ruthenborg, R. J., Ban, J. J., Wazir, A., Takeda, N. & Kim, J. W. Regulation of wound healing and fibrosis by hypoxia and hypoxia-inducible factor-1. Mol. Cells 37, 637–643 (2014). [10.14348/molcells.2014.0150]

41. Cialdai, F., Risaliti, C. & Monici, M. Role of fibroblasts in wound healing and tissue remodeling on Earth and in space. Front. Bioeng. Biotechnol. 10, 958381 (2022). [10.3389/fbioe.2022.958381]

42. Han, C., Barakat, M. & DiPietro, L. A. Angiogenesis in wound repair: too much of a good thing? Cold Spring Harb. Perspect. Biol. 14, a041225 (2022). [10.1101/cshperspect.a041225]

43. Choi, J., Hong, G., Kwon, T. & Lim, J. O. Fabrication of oxygen releasing scaffold by embedding H2O2-PLGA microspheres into alginate-based hydrogel sponge and its application for wound healing. Appl. Sci. 8, 1492 (2018). [10.3390/app8091492]

44. Rafique, M. et al. Insight on oxygen-supplying biomaterials used to enhance cell survival, retention, and engraftment for tissue repair. Biomedicines 11, 1592 (2023). [10.3390/biomedicines11061592]

45. Abdi, S. I. H., Choi, J. Y. & Lau, H. C. et al. Controlled release of oxygen from PLGA– alginate layered matrix and its in vitro characterization on the viability of muscle cells under hypoxic environment. Tissue Eng. Regen. Med. 10, 131–138 (2013). [10.1007/s13770-013-0391-7]

46. Guo, J. et al. Implications of pH and ionic environment in chronic diabetic wounds: an overlooked perspective. Clin. Cosmet. Investig. Dermatol. 17, 2669–2686 (2024). [10.2147/CCID.S485138]

47. He, Y., Wang, W. & Ding, J. Effects of L-lactic acid and D,L-lactic acid on viability and osteogenic differentiation of mesenchymal stem cells. Chin. Sci. Bull. 58, 2404–2411 (2013). [10.1007/s11434-013-5798-y]

48. Monfoulet, L. E. et al. The pH in the microenvironment of human mesenchymal stem cells is a critical factor for optimal osteogenesis in tissue-engineered constructs. Tissue Eng. Part A 20, 1827–1840 (2014). [10.1089/ten.TEA.2013.0500]

49. Thacker, M., Chen, Y. N., Lin, C. P. & Lin, F. H. Nitrogen-doped titanium dioxide mixed with calcium peroxide and methylcellulose for dental bleaching under visible light activation. Int. J. Mol. Sci. 22, 3759 (2021). [10.3390/ijms22073759]

50. Bankar, N. et al. Antimicrobial and antibiotic-potentiating effect of calcium peroxide nanoparticles on oral bacterial biofilms. NPJ Biofilms Microbiomes 10, 106 (2024). [10.1038/s41522-024-00569-7]

51. Huang, L., Chen, X., Yang, X., Zhang, Y. & Qiu, X. GelMA-based hydrogel biomaterial scaffold: a versatile platform for regenerative endodontics. J. Biomed. Mater. Res. B Appl. Biomater. 112, e35412 (2024). [10.1002/jbm.b.35412]

52. Ansari, S. et al. RGD-modified alginate–GelMA hydrogel sheet containing gingival mesenchymal stem cells: a unique platform for wound healing and soft tissue regeneration. ACS Biomater. Sci. Eng. 7, 3774–3782 (2021). [10.1021/acsbiomaterials.0c01571]

53. United States Environmental Protection Agency. Calcium peroxide fact sheet – registered uses and regulatory status. Office of Pesticide Programs, US EPA (2022).

54. International Organization for Standardization. ISO 10993-1:2018. Biological evaluation of medical devices – Part 1: Evaluation and testing. ISO (2018).

55. Augustine, R. et al. Oxygen-generating scaffolds: one step closer to the clinical translation of tissue engineered products. Chem. Eng. J. 455, 140783 (2023). [10.1016/j.cej.2022.140783]

56. Lu, P., Ruan, D. & Huang, M. et al. Harnessing the potential of hydrogels for advanced therapeutic applications: current achievements and future directions. Signal Transduct. Target. Ther. 9, 166 (2024). [10.1038/s41392-024-01852-x]

57. Yang, M. et al. Hydrogel microspheres as versatile platforms for biomedical research: design, properties, and applications. MedComm 6, e70423 (2025). [10.1002/mco2.70423]

